# Dissecting the steps in early Simian Immunodeficiency Virus dissemination following mucosal and intravenous infection of rhesus macaques

**DOI:** 10.1101/2025.03.24.644849

**Authors:** Steffen S. Docken, Agatha Macairan, Timothy E. Schlub, Christine M. Fennessey, Benjamin Varco-Merth, Louis J. Picker, Afam A. Okoye, Taina T. Immonen, Deborah Cromer, Brandon F. Keele, Miles P. Davenport

## Abstract

In cases of HIV transmission, the typical delay from exposure to detectable viremia is approximately one week. This delay from exposure to viremia suggests that during initial expansion of virus from a limited number of founder lineages, there exists a period of low infected cell population. It is during this period of low infected cell population that the virus may be more vulnerable to clearance via primed immune responses targeting infected cells (e.g., antibody-dependent cellular cytotoxicity (ADCC) or CD8^+^ T cell killing/suppression). Potential future prophylactic harnessing of these immune mechanisms for early virally infected cell clearance will rely on an understanding of the earliest stages of viral replication and dissemination. The factors that dictate the rate of early viral spread, termed the ‘dissemination bottleneck’ could include target-cell-mediated effects, the anatomical microenvironment, or the organ of the first infected cells. In this study, we use the barcoded Simian Immunodeficiency Virus (SIV) infection model to assess the contribution of various anatomical and cellular mechanisms to the SIV dissemination bottleneck. Viral, cellular, and anatomically-mediated heterogeneity in viral replication each introduce a degree of variability into the early phases of viral spread, and when multiple founder lineages are present, this variability in early growth results in a large distribution in lineage sizes. Therefore, we use a comparison of viral lineage size variability across multiple experimental SIV infection models to examine the relative contribution to the overall dissemination bottleneck of viral-mediated stochasticity of cellular infection (e.g., integration site), infected cell phenotype (e.g., activation state), anatomical variability, and initial viral spread within the genital tract. We estimate that inherent heterogeneity in viral production by infected cells corresponds to 23 to 44% of the dissemination bottleneck, but the majority (56 to 77%) arises from anatomical heterogeneity (presumably heterogeneity in how conducive local microenvironments are to viral replication).

**Author Summary:** A brief window exists immediately following HIV transmission where low initial levels of virus and infected cells may be susceptible to immune clearance. A better understanding of the bottlenecks encountered by the virus during this window is necessary when designing therapies to clear HIV during this early period. We used a barcoded Simian Immunodeficiency Virus (SIV) infection model to track the early dissemination of multiple viral lineages after mucosal and intravenous inoculation. We observed up to 10^5^-fold differences in lineage size between transmitted barcodes within a single animal two weeks after infection, suggesting very different trajectories of virial growth. By comparing lineage size diversity after mucosal and intravenous transmission and in vitro replication, we determined the contribution of anatomical and cellular mechanisms to the relative growth of different clonotypes. Although we expected that the processes of local dissemination may lead to greater lineage size diversity after mucosal transmission, this was not the case, and we saw no difference compared to intravenous transmission. We found that around a quarter of the diversity in clonotype size could be attributed to early cellular infection events, with the remainder likely attributable to differences in clonotype specific establishment and dissemination in vivo.

## Introduction

It is known that HIV transmission involves a stringent bottleneck, meaning that only a small proportion of exposure events lead to transmission [1–4]. Furthermore, when transmission does occur, infection appears to be established by only a single founding virion in ∼80% of sexual transmission cases [5–8]. As a result, many studies have focused on understanding the first lines of defense such as the mucosal barrier [7, 9, 10]. Additionally, neutralizing antibodies [11, 12] may play a role in reducing the initial entry of virus if given passively or vaccine-induced, thereby blocking infection altogether. However, following transmission, there is also a substantial delay from the time of exposure to the detection of widespread systemic replication and viremia [13, 14]. This delay results from the expansion of virus from a single or very few infected cells, which can occur in various sites depending on the initial exposure route of the virus and its subsequent spread. During the early phases of this expansion, the limited amount of virus with few infected cells is particularly vulnerable to immune control or clearance via mechanisms such as antibody-dependent cellular cytotoxicity (ADCC) [15, 16] or CD8^+^ T cell responses to infection [17, 18]. Thus, in addition to the entry bottleneck for HIV infection we might also consider an early dissemination bottleneck, during which the small population of infected cells remains particularly vulnerable to immune targeting. We could then consider how different vaccines or immune interventions may be focused to eliminate or further delay viral spread [11, 18]. Thus, dissecting the mechanisms underlying the dissemination bottleneck will be critical to application of immunoprevention strategies.

In the case of sexual transmission of HIV, the virus is thought to initially traverse the mucosal barrier (transmission bottleneck) [7, 9], before infecting local target cells (predominantly CD4^+^ T cells within the genital tract), and subsequently spreading more widely [19]. Studies of mucosal transmission in non-human primate SIV infection models aim to mimic this process of spread from entry to viremia [20].

Non-human primate studies using a barcoded SIVmac239 to study early events following intravaginal (mucosal) SIV inoculation of rhesus macaques suggest that even when only a few virions establish infection, they have a wide diversity of subsequent dissemination kinetics [21]. In these animals, we observed large heterogeneity in lineage size both within the female genital tract during early infection, as well as in subsequent sites of dissemination (e.g., in plasma, lymph nodes, spleen, etc.), and in plasma viraemia. This heterogeneity in lineage size is indicative of stochastic delays in viral growth and spread early in infection (i.e., a dissemination bottleneck).

Even with these non-human primate models, direct analysis of early kinetics of SIV/HIV dissemination is difficult. Potential mechanisms contributing to the dissemination bottleneck range in physical scale from molecular differences in viral strains (e.g., viral fitness), to the cellular phenotype and viral production by the first infected cell, to the microenvironment or organ in which the first infected cell is located and the distance to lymphoid sites. For example, studies of infection with a combination of two lineages varying by a single escape mutation suggest that in the absence of immune targeting, more fit wild-type virus can double in relative frequency every 2 days [22, 23]. Additionally, at the cellular level, image analysis of infected cells demonstrates that the viral production per cell can vary by at least 10-fold (although this could be much higher, due to the limit of detection of low viral production) [24]. Lineages established by infection of high-producing cells would be expected to grow faster and to progress to systemic replication more rapidly than lineages established by low producing cells. At a slightly larger scale, characteristics of the microenvironment of the first infected cells, such as concentration and activation level of the available target cells could affect the rates of both viral production and spread within the tissue. From the perspective of immune control of dissemination, delays in early viral spread may provide a larger window in which mechanisms such as CD8^+^ T cell killing or ADCC may act with greater efficacy on a smaller infected target cell population.

In this study, we use a barcoded SIV virus model to better understand the variability in the trajectories of viral dissemination that can be observed following infection of non-human primates with a population of phenotypically identical viruses. We observed greater than a 100,000-fold difference in viral clonotype (barcode) sizes during early infection in some animals, even when each clonotype infection is thought to be initiated by a single virion. We compare this diversity in clonotype sizes with the diversity seen after single or multiple rounds of in vitro infection of either primary CD4^+^ lymphocytes or a susceptible cell line. We estimate that between 23 and 44% of the variability in clonotype size can be accounted for by heterogeneity in the amount of virus produced by individual cells during very early infection, with aspects of anatomical variability (e.g., potentially the microenvironment of some of the first infected cells) accounting for the remaining 56 to 77% of the observed variability in lineage size. This suggests a potential window of vulnerability for the virus during low-level replication in the local microenvironment, prior to more widespread dissemination.

## Results

### SIV dissemination and clonotype size diversity after intravaginal versus intravenous infection

The pathway of the virus from the site of exposure to systemic infection after mucosal inoculation involves multiple steps, including transiting the mucosal surface, early local spread in the genital tract and then to local lymphoid tissue, and later dissemination. In contrast, intravenous inoculation bypasses the need for mucosal transit and local spread but still requires systemic establishment and spread. Therefore, comparison of the early viral dynamics of these two infection models provides insight into the impact of mucosal tissue on the dissemination bottleneck.

In a previous study, we have observed substantial heterogeneity in viral clonotype size after intravaginal infection of rhesus macaques with a barcoded SIV [21]. In this previous study, animals were infected via intravaginal (mucosal) inoculation (n = 5) using a pool of equivalent titers of 10 replication competent barcoded SIV strains (SIVmac239X), of which 9 were deemed suitable for analysis in our current study (see Animal experiments in Materials and Methods). The inoculating challenge contained 10^6^ infection units (IU). Assessment of early transmission events after mucosal inoculation suggested that transmission occurred in a small number of sites in the vagina [21]. Additionally, sequencing of subsequent plasma viremia revealed an average of only 5.6 barcodes established infection following mucosal inoculation (range: 2 to 8 barcodes), suggesting that most established barcodes were ‘founded’ by only one or two infecting virions. These mucosally infected animals were necropsied (between day 6 and 14) prior to peak viral load, with viral loads ranging from 2.0 × 10^3^ to 1.4 × 10^7^ copies/ml on the day of necropsy [21].

For the intravenous inoculation study, we used animals infected intravenously with a barcoded virus with high barcode diversity (SIVmac239M), which has over 10,000 distinct barcoded lineages within the stock [25]. The intravenously inoculated animals were challenged with 100, 200, 500, or 2200 IU of virus, and plasma viral RNA was sequenced 8, 11, 12, or 15 dpi [26, 27]. The peak viral load was similar across the four inoculation sizes (geometric mean of 3.2 × 10^7^ copies/ml, range 3.3 × 10^5^ to 2.8 × 10^8^ copies/ml), but timing of peak viral load depended on inoculation size (mean of day 18, 13, 12, and 12 for inoculations of 100, 200, 500, and 2200 IU respectively). The number of barcodes detected in plasma was highly correlated with inoculation size (S1 Fig.) and ranged from 4 to 387 (S1 Table). However, since the inoculating virus contained over 10,000 barcodes [25] and less than 4.2% of the barcodes establish infection in any one animal, we assume that each barcode represented the progeny of only one or a very small number of infecting virions.

Despite most barcode lineages were founded by a single infection event, we observed a high diversity in the fractional contribution of individual barcoded lineages to the total plasma viral load early after infection (Fig 1B-E).

**Fig 1:**
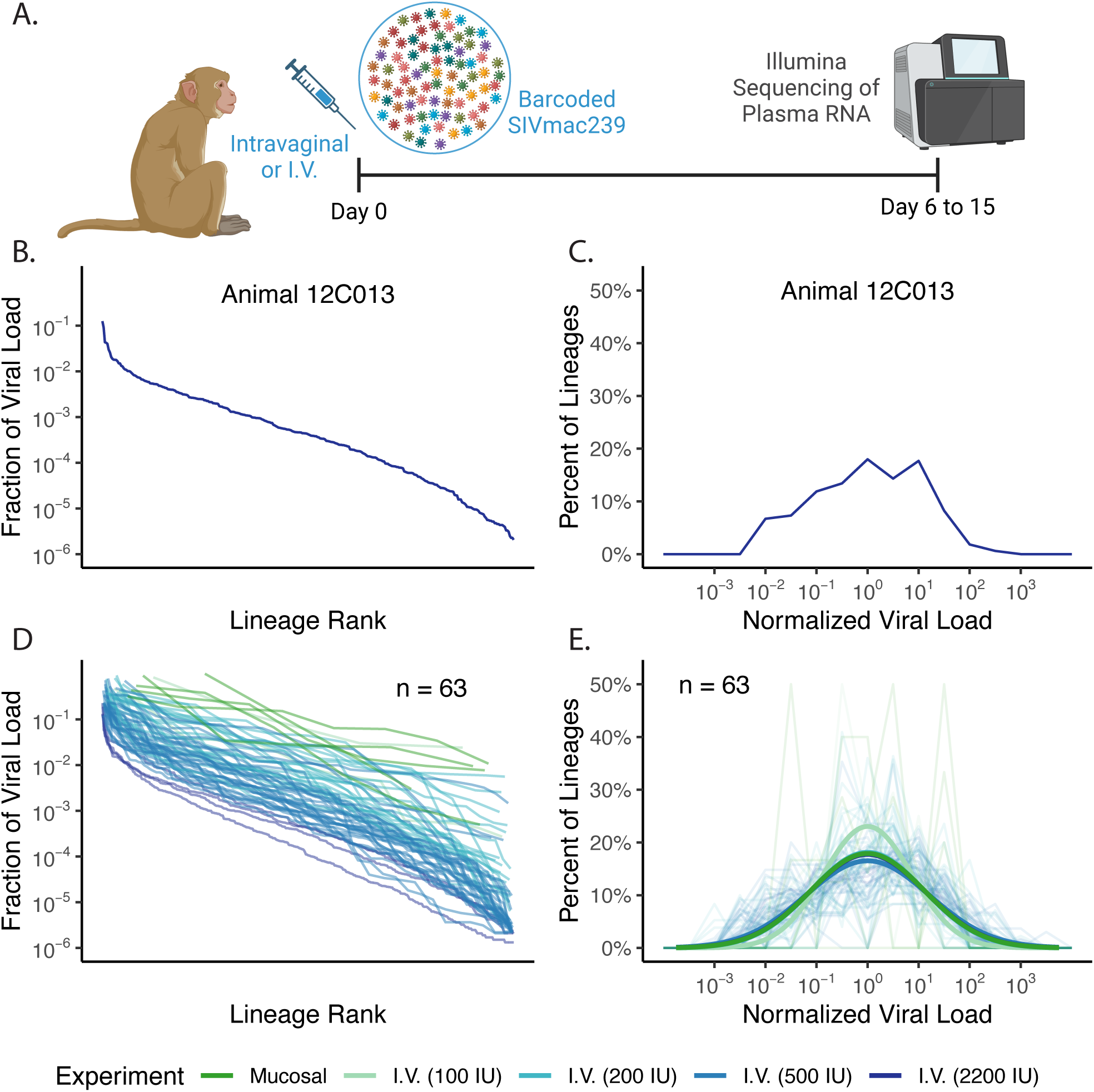
Viral lineage size heterogeneity in vivo following intravaginal versus intravenous inoculation. (A) Schematic of experimental protocol for rhesus macaque studies with intravaginal (mucosal) or intravenous inoculation. Created in https://BioRender.com. (B and C) Viral lineage rank abundance curve (B) and corresponding empirical distribution (C) for a typical animal inoculated intravenously with 2200 IU. In panel (B), the horizontal-axis shows the rank order of barcode size from largest to smallest, and the vertical-axis shows the fraction of the total viral load for the relevant barcode (i.e., the proportion of read-counts from that barcode). (D and E) Viral lineage rank abundance curves (D) and empirical distributions of lineage (E; faded lines) for all mucosally (dark green) and intravenously inoculated animals (shades of light green to dark blue indicate inoculation size). Empirical distributions for each animal were generated by normalizing lineage sizes by the geometric mean and binning lineages by 0.5 log10 intervals. Dark solid lines in (E) illustrate best fit log-normal distributions for mucosal inoculation and each intravenous inoculation dose.

Since for both the mucosal and intravenous transmission routes, we believe infection was initiated by only one or two infecting virions per barcode clonotype, the diversity of lineage proportion sizes observed in plasma after infection is likely attributed to the effects of subsequent processes involved in viral dissemination. Therefore, comparison of the distributions in lineage proportions between animals infected mucosally or intravenously will indicate the impact of infection route on early dissemination. An example of the observed distribution in the sizes of each barcoded lineage for a typical intravenously inoculated animal is shown in Fig 1B and C, as a rank abundance curve (Fig 1B) and empirical distribution (Fig 1C). Similar data across all mucosally or intravenously inoculated animals are shown in Fig 1D (rank abundance curves) and the faded lines in Fig 1E (empirical distributions). We observe a log-normal distribution in barcoded viral lineage sizes within individual animals (Fig 1C and E), and we assume that this heterogeneity in lineage sizes within an animal is a proxy for the stringency of the relevant dissemination bottlenecks that the virus has needed to navigate in the two experimental systems. Therefore, we seek to quantify this lineage size heterogeneity and use the variance of log_10_ lineage size within an animal as a measure of heterogeneity. (For simplicity, going forward we will just refer to lineage size variance, with the log_10_ implied.) We calculated the mean lineage size variance as 1.24 ± 0.54 (mean ± standard error) in the mucosally infected animals and 1.33 ± 0.07 in the intravenously infected animals. The intravenously infected animals had a slightly higher but not significantly different variance than intravaginally inoculated animals (see S1 Table for breakdown by inoculation size). Based on log-normal distributions with these variances, we estimate that 63% of infecting lineages would be within 10-fold of the geometric mean lineage size in the intravaginally inoculated animals, and 61% of lineages would be expected within 10-fold of the geometric mean lineage size in the intravenously inoculated animals (see Mathematical modeling and statistical analysis in Materials and Methods). This suggests that despite the differences in infection routes and the need for local dissemination after mucosal infection, the bottlenecks encountered and subsequent diversity of clonotype size are very similar after intravenous versus mucosal exposure. In other words, we do not see evidence of local spread within the genital tract contributing additional or more stringent dissemination bottlenecks than those encountered in other initial sites of infection (i.e., tissues infected immediately following intravenous inoculation).

### Evidence for lineage size diversity in vitro

The results above suggest that the large clonal size heterogeneity observed after mucosal inoculation is independent of the effects of local dissemination, since it is also observed after intravenous infection. Instead, downstream events in viral dissemination common to both routes of inoculation seem likely to be the major contributors to diversity. These include differences in levels of viral production by the first infected cell(s), variability in the number of secondary cells infected by the first infected cell, and differences in rate of spread due to presence in distinct anatomical niches. Although it is difficult to directly assess the impact of distinct anatomical niches on viral dissemination, it is possible to study cellular viral production and early viral spread in vitro and assess whether these might be important in determining lineage size diversity. Therefore, we sought to measure the heterogeneity in viral production following in vitro infection of stimulated primary CD4^+^ T cells from rhesus macaques with barcoded virus (SIVmac239V67M; for details, see In vitro cultures in Materials and Methods). We used a low multiplicity of infection (MOI = 0.003) so that each barcode would infect only a single cell. To restrict virus to a single round of replication cells were treated with anti-retrovirals (maraviroc and emtricitabine – FTC starting at 18 and 24 hours post infection, respectively), and to block virion attachment to cells after production, anti-CD4 antibodies were used (starting at 18 hours post infection). We monitored viral production by individual infected cells between 24 to 48 hours post infection (see Materials and Methods for details).

Collection and viral quantification of the supernatant of three replicate cultures revealed that the geometric mean of virus production between 24 and 48 hours post infection was 1.5 × 10^6^ viral copies (range 1.4 × 10^6^ to 1.7 × 10^6^ copies). Sequencing identified between 1,192 and 1,305 different barcodes from each well, which corresponds to less than 5% of the barcodes in the stock. Because of the high diversity of barcodes and low MOI, this indicates each barcode likely corresponds to a single infected cell.

In the sequencing of each well of primary cells, we again observed a roughly log-normal distribution of barcodes with a high number of copies (main panel of Fig 2B), as well as a large tail of barcodes detected at low frequency (left side of inset in Fig 2B). The log-normal distribution appeared consistent with the distribution observed in vivo. However, it was not clear whether the ‘tail’ of low frequency barcodes represented an artefact such as carry over of inoculum, sequencing error [25], some low level of abortive infection, or if this tail is excluded in vivo due to active clearance mechanisms. To characterize the distribution of viral production by primary cells, we fit a distribution composed of power-law and log-normal components corresponding to the low frequency tail and the high frequency barcodes, respectively (dark lines in Fig 2B and its inset; see Mathematical modeling and statistical analysis in Materials and Methods). To analyze the clonotype size diversity after single round infection in vitro, we ignored the low producing lineages in the tail (we note that this is a conservative assumption, since it leads to a narrower lineage size heterogeneity) and focused on the log-normal distribution of the barcodes with a high number of copies. From our fitted distribution, we estimated a mean lineage size variance for these high producing primary cells of 0.31 ± 0.01. This is a substantially narrower distribution than in the intravaginally or intravenously inoculated animals and suggests that for 93% of stimulated primary cells, viral production is within 10-fold of the geometric mean production (see Materials and Methods for details).

**Fig 2:**
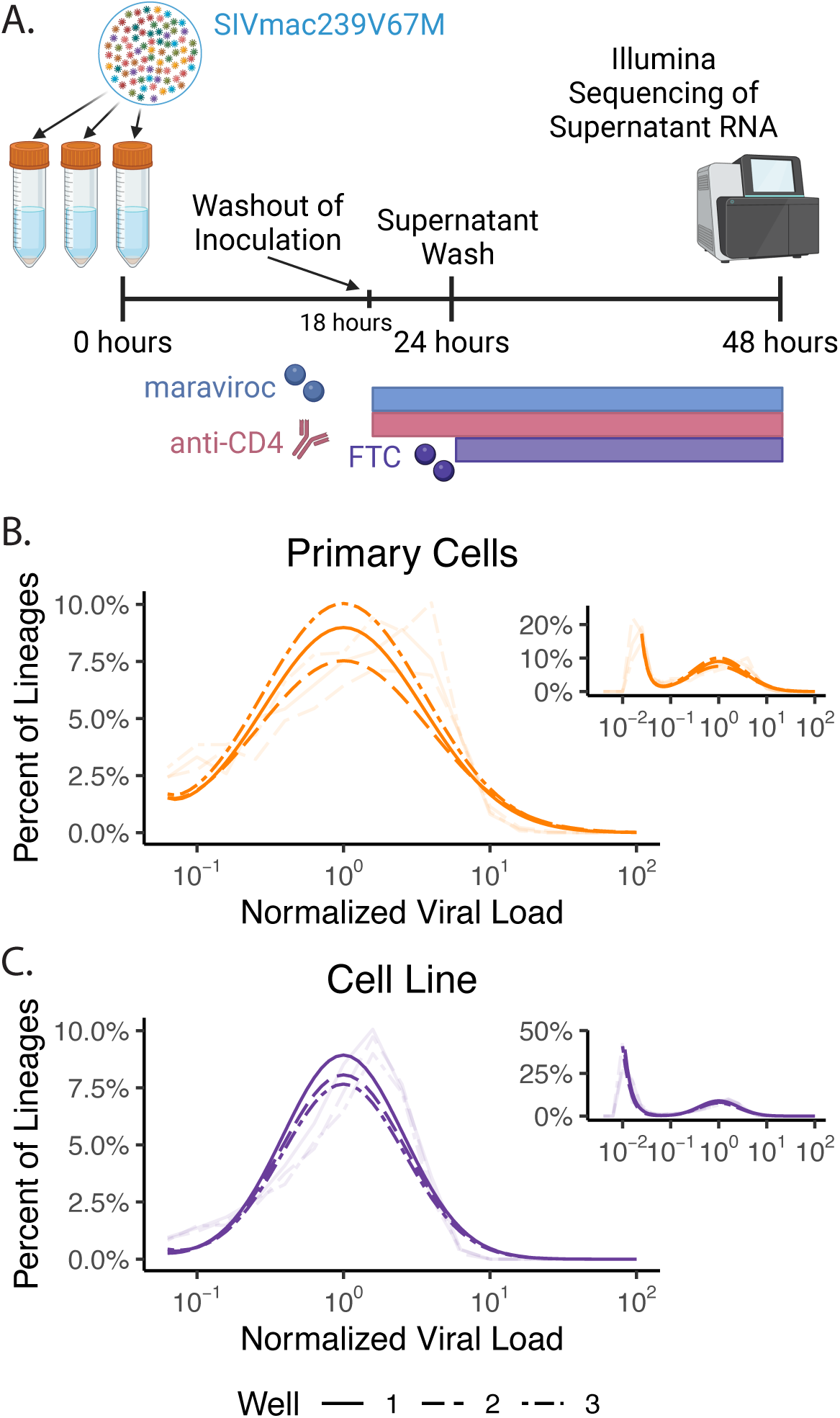
Lineage size heterogeneity in vitro. (A) Protocol for in vitro infection of stimulated primary cells or SupT1-R5 cells with SIVmac239V67M. Inoculation washed out after ∼18 hours, supernatant washed at 24 hours, and supernatant collected at 48 hours for barcode sequencing. Wells were treated with maraviroc and anti-CD4 antibody starting at 18 hours post inoculation and FTC starting at 24 hours to block infection of new cells. Created in https://BioRender.com. (B-C) Empirical distribution (faded lines) and best fit distributions (dark lines) of lineage size (log10-scale) in each of the three stimulated primary cell (B, orange lines), and SupT1-R5 wells (C, purple lines).

It is clear in the analysis above that there is a significant variability in viral production from individual cells following a single round of infection in primary cells. However, it is not clear whether this arises from cell-mediated effects such as phenotypic heterogeneity, or from virally-mediated effects (potentially integration site heterogeneity). Therefore, we repeated the in vitro infection using the SupT-R5 cell line, so as to minimize phenotypic heterogeneity among the target cells. Between 24 and 48 hours post infection, SupT-R5 cells in the three wells produced a geometric mean of 1.3 × 10^6^ viral copies (range 1.2 × 10^6^ to 1.4 × 10^6^ copies), and sequencing of the supernatant detected between 2,932 and 3,023 different barcodes per well (10% or less of the inoculating barcodes). Again, we fit the distribution consisting of power-law tail and log-normal components to the barcode frequency data (Fig 2C).

The mean lineage size variance among the high producing cells after infection of SupT-R5 cells was only 0.173 ± 0.002, which is significantly lower than in the more phenotypically diverse stimulated primary cells (p = 0.008). This significant decrease in viral production heterogeneity suggests that the diversity in infected cell phenotype in primary cells may have a significant impact on the amount of virus produced by the cell. However, the mean lineage size variance of 0.17 (expectation of 98% of lineages within 10-fold of the geometric mean; Mathematical modeling and statistical analysis in Materials and Methods) in the cell line indicates that even among notionally homogeneous cells, heterogeneity in infection outcome (perhaps as a result of factors such as integration site) causes variability in the amount of virus produced by different cells.

Potential alternative explanations for the observed variability in barcode sizes in the in vitro infection models are differences in viral fitness across barcoded lineages or PCR amplification bias artificially inflating lineage size heterogeneity. However, previous studies have demonstrated that the barcoded viruses used here are functionally similar [25, 26, 28]. Alternatively, if all lineages were actually similar in size and the observed heterogeneity was merely due to PCR amplification bias, PCR amplification of a secondary sample would generate a complete reshuffling of the lineage size rankings. Therefore, we compared measured barcode frequency at days 8 and 14 in four of the intravenously inoculated animals, and we observed an extremely high correlation in barcode frequency between these two time points (p < 10^-24^; S1 Text), supporting the validity of the observed biological variation in lineage size. Furthermore, we performed PCR amplification on highly diluted viral stock, such that individual barcodes are expected to be present at only a single copy. We found a low variance in the amplification of individual barcode templates that was insufficient to explain the observed heterogeneity in single cell production (mean variance of 0.013; see S1 Text for further details).

### Sources of lineage size heterogeneity during SIV dissemination

To assess the contribution of each potential source of variability in the establishment of early infection to the dissemination bottleneck, we compared lineage size variance estimates across the in vivo and in vitro experimental setups (described in the first two sections of the Results). To allow for direct comparison of the contributions of different sources of variability, we express the heterogeneity in lineage size as a proportion of the heterogeneity seen after intravenous inoculation.

The diversity observed after infection of the SupT1-R5 cell line demonstrates that a significant proportion of the final lineage size heterogeneity occurs even when identical virions (apart from the barcode) infect a phenotypically homogenous cell line (estimated 12% of the heterogeneity following intravenous inoculation, Fig 3A and B; Estimation of proportional breakdown of dissemination bottleneck in Materials and Methods). This suggests that random viral factors such as integration site may affect subsequent viral production levels. Infection of stimulated primary cells leads to significantly greater heterogeneity (an additional 11% of heterogeneity following intravenous inoculation, p = 0.008 compared to infection of cell line). This suggests that infection of a more heterogenous cell population may lead to a greater heterogeneity in viral production levels.

**Fig 3:**
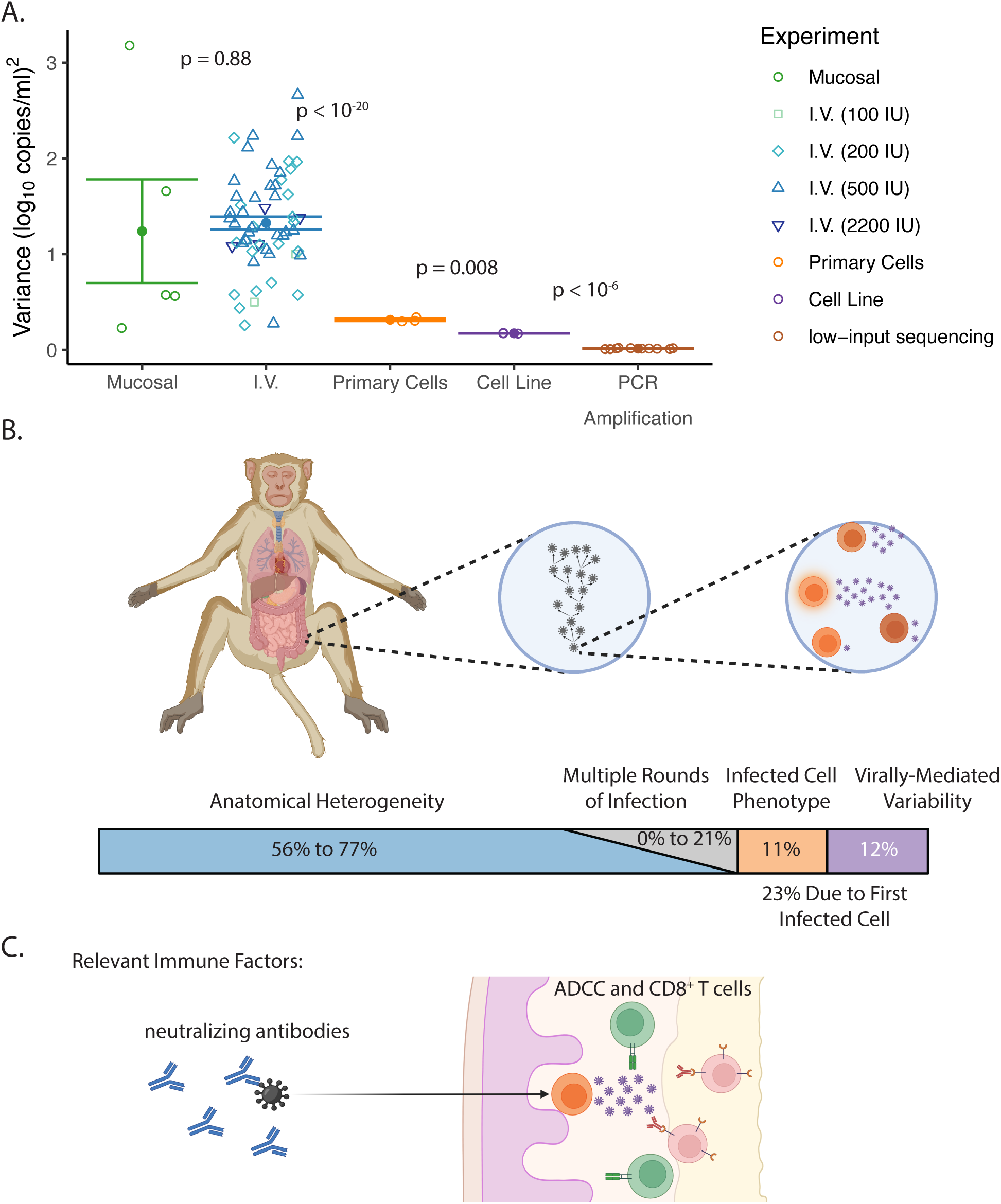
Lineage heterogeneity across experimental set ups. (A) Variance in log10 lineage size for all mucosally inoculated animals (green circles), intravenously inoculated animals (blue; shade and shape indicates inoculation size), in vitro primary cell wells (orange circles), SupT1-R5 wells (purple circles), and PCR amplification of highly diluted viral stock (brown circles) is plotted. Solid points and error bars indicate mean and standard error of variance in each experimental set up. (B) The partitioned bar indicates the percentages of the dissemination bottleneck corresponding to (from left to right) different anatomical niches of replication, accumulated heterogeneity over multiple rounds of replication (grey), and heterogeneity in production by the first infected cell due to cell phenotype and virally-mediated variability. Percentages of the total dissemination bottleneck were estimated based on comparisons to lineage size variance in intravenously inoculated animals following background subtraction of PCR amplification variance (see Estimation of proportional breakdown of dissemination bottleneck in Materials and Methods). (C) Schematic representation of where/when various immune factors may be able to eliminate the virus during the transmission process (i.e., targeting either the transmitted virus or the low viral loads of early infection), as previously described in Picker et al. 2012 and 2023 [17, 18]. Components of (B) and (C) created in https://BioRender.com.

The viral size heterogeneity discussed above focusses on the first round of infection only (which was then arrested in vitro by treatment). However, modeling suggests that applying a similar distribution in viral production over several rounds of infection can further amplify this heterogeneity [29]. To examine this, we analyzed lineage size variance after multiple days of infection in vitro (in the absence of anti-retroviral treatment) and compared to the variance following a single cycle of production. We find that the heterogeneity of lineage size more than doubled over seven days of viral spread in the SupT1-R5 cell line (mean variance of 0.45 ± 0.02 (log_10_ copies/ml)^2^; see S2 Text). This increase in heterogeneity was not observed in infection of stimulated primary cells, likely because of a limited spread of virus in these cultures (as evidenced by reduced viral growth beyond 4 days of culture). The mean lineage size variance in these cultures at 4 days was 0.28 ± 0.01 (log_10_ copies/ml)^2^ (see S2 Text) vs. 0.31 ± 0.01 (log_10_ copies/ml)^2^ from single cycle replication. Based on the observed increase in lineage size variance in the cell line cultures compared to the total variance in intravenously inoculated animals, we estimate that the cumulative effects of viral production heterogeneity across subsequent rounds of infection may cause up to an additional 21% of the total dissemination bottleneck (Fig 3B, S2 Text).

The estimates above suggest that lineage size variance after intravenous inoculation is still at least twice the variance that might be predicted from the in vitro infection model. A proportion of this may simply reflect the larger cellular heterogeneity in vivo (i.e., there may be more phenotypic diversity found in the whole body than reflected in stimulated primary cells). The clustering of phenotypically similar cells in vivo may further amplify differences in cell growth. That is, a virus infecting a site with a high density of activated cells may rapidly spread from one highly productive cell to another (compared to a cell in a more quiescent or mixed environment). The effect of local microenvironmental differences would not be captured in vitro, since cells with different phenotype and activation state are assumed to be relatively well mixed in the culture well.

Previous studies have described the process of local spread after mucosal inoculation [20, 21, 30]. Surprisingly, we saw no difference in lineage size variance comparing intravenous and mucosal inoculation (Fig 3A). Thus, whatever additional processes occur after mucosal infection do not appear to add to lineage size heterogeneity, and hence the dissemination bottleneck.

## Discussion

Barcoded viruses provide the opportunity to study viral growth trajectories of multiple independent infection events in the same animal. We analyzed historic data on individual barcode frequency following intravenous (n = 58 [26, 27]) or mucosal infection (n = 5 [21]) of non-human primates. We observed that individual lineages varied up to 100,000-fold in their copy number in serum as early as 15 days after inoculation. This clonotype size diversity suggests that bottlenecks in viral dissemination can act in a highly variable way to limit or facilitate the spread of virus after successful infection. To dissect the possible mechanisms of heterogeneity in viral establishment and growth we turned to an in vitro infection model also using a barcoded virus to further clarify the earliest viral growth trajectories. We demonstrate that nearly a quarter of total in vivo clonotype size diversity may arise from stochastic events occurring during infection of the first cell, with further diversity accumulating over the first few infection cycles (Fig 3B). Compared to the in vivo SIV-infected rhesus macaque model, which necessarily includes anatomical complexity, we found lineage size variance was reduced by over 50% in the in vitro models. Therefore, we estimate that over 50% of the dissemination bottleneck is unmeasured and likely arises from diversity in anatomical niches.

The barcoded SIV viruses used in this study are thought to be phenotypically identical [25, 26, 28], as indicated by the observation of dominance of different clonotypes between animals as well as the parallel growth of barcodes in plasma within the same animal (S1 Text). This variation in dominant barcodes is consistent with a model of stochastic differences in viral spread early in infection (when small infected cell numbers and localized infection may amplify differences in viral production between cells and microenvironments), followed by similar growth of barcodes after wider dissemination of infection. Although it is not clear at what point viral growth rate might become consistent between barcodes, we can estimate the effective differences in timing between when the largest and smallest clonotypes reach this threshold. For example, the difference in clonotype size between the 10^th^ and 90^th^ centiles of clonotypes in vivo is around 1700-fold. Since the average viral growth rate across intravenously inoculated animals in this study is around 1.9 per day (corresponding to a 6.8-fold daily increase in viral load), a 1700-fold difference equates to around a 4-day delay to the start of consistent exponential growth. This suggests that some clonotypes may spend an extended period at lower viral levels, creating an extended window for potential immune control or natural extinction of a proportion of viral lineages (Fig 3C). In the context of natural infection with a single founder virus (or low-dose infection in SIV), this suggests that individual infection events may be vulnerable to early immune control for quite variable periods of time. This may explain, for example, why CMV-vectored vaccines are able to reduce the establishment of infection by up to 59% (due to immune control of a proportion of early infection events), but if infection is established in vaccinated animals, it follows an apparently normal trajectory (once viral replication becomes widespread) [18, 31–33].

The large difference in clonotype diversity between in vivo and in vitro studies indicates that there is a significant influence of unmeasured heterogeneity in lineage size variance. One mechanism may be the diversity of anatomical niches in which the virus becomes established. These niches might vary in both the phenotype and density of potential target cells, creating either ‘deserts’ or ‘jackpots’ for founder virus establishment and growth. Since we observed similar lineage size variances following both mucosal and intravenous inoculation routes, this suggests that the wide in vivo lineage size variation is not dependent on mucosal entry/pathways or dissemination from mucosal sites.

Our study uses barcoded, functionally identical viruses to simultaneously study multiple viral lineages in vivo and in vitro. Although this infection model is incredibly insightful, there are a few of limitations with respect to our study. Firstly, the scope of our examination of clonotype size diversity is confined to that caused by host-mediated bottlenecks, rather than due to differences in viral phenotype, due to the lack of viral phenotypic variability in our experiments. Yet, it is known that diversity in viral phenotype also affects cellular infectivity and production rates [34–37], and thereby likely also affects local spread within organs or microenvironments; hence impacting the dissemination bottleneck. Therefore, our estimates are likely an underestimation of the total viral heterogeneity in lineage size during primary infection if infection begins with two or more phenotypically and replicatively distinct lineages. Secondly, we note that the mucosally inoculated animals were inoculated with fewer barcodes (9 functionally similar barcodes) than the intravenously inoculated animals. This may have resulted in some barcode lineages being initiated by multiple founder virions and could therefore have impacted our lineage size variance estimates. However, we anticipate this had only a limited effect, and that most barcode lineages were initiated by only one or two infecting virions. This is based on the fact that an average of only 5.6 barcodes were detected per animal, out of a possible 9 barcodes. Similarly, in our in vitro studies infecting with barcoded virus, which represent a novel approach to quantifying the production of virus from individual infected cells, it is statistically possible a small proportion of viral barcodes were be produced by more than one infected cell.

An important finding from this study is the diversity in production rates of virus from a single cell. Among the stimulated primary cells, the 10% most productive cells were responsible for producing somewhere between around 30 - 40% of virions. Notably, this may be an underestimate of the variability in viral production due to the fact that the primary cells in our analysis received a uniform stimulus (PHA stimulation), as opposed to the diverse range of stimulating signals cells receive in vivo. Therefore, our in vitro experiments (used to calculate this variability in viral production) likely represent only a subset of the full phenotypic diversity of CD4^+^ T cells in vivo. Our finding of a wide distribution of viral production following SIV infection of single cells is very consistent with previous observations of viral production of HIV ex vivo [24, 38]. Hataye et al. stimulated limiting dilutions of latently infected cells from people living with HIV and monitored if viral production occurred over the following 12 days [38]. Across the 42 wells with detected production, the variance in viral load on the first day of viral detection was 0.69 (log_10_ copies)^2^. This suggests that production by 77% of reactivating latently infected cells falls within 10-fold of the geometric mean viral load (Mathematical modeling and statistical analysis in Materials and Methods). Taken together, our study and the results of Hataye et. al. suggest that only a small subset of infected cells produce the bulk of free virus and are the key drivers of ongoing replication. By contrast, a significant proportion of infected cells may produce insufficient virus for ongoing propagation of infection (depending on the local microenvironment). Identification of the cell phenotypes and viral factors that define this elite subset of high-producing infected cells may allow us to target this subset to reduce the probability of establishing infection.

Studying the early events in HIV/SIV infection is challenging, as it requires looking for a ‘needle in a haystack’ to identify the first infection events [20, 21, 30]. The use of barcoded virus allows us to simultaneously study multiple infection trajectories in the same animal or in vitro cultures. Our analysis demonstrates that early infection plays out quite differently for individual lineages even in a naïve animal, and this may also have important implications for viral susceptibility to immune control during early infection. Further work is required to identify the cellular and anatomical determinants of the rate of viral spread, as well as to determine the implications of this for potential immune control in early infection.

## Materials and Methods

### Experimental protocols

#### Animal experiments

The experimental protocol for the intravaginally inoculated animals was previously described in Deleage et al. [21]. In that study, fifteen Indian-origin rhesus macaques were intravaginally challenged with 10^6^ IU (TZM-bl) of SIVmac239X (10 genetically distinct viral lineages described previously [28]) and serially necropsied between 3 and 14 days post infection (dpi) [21]. Five of these animals had detectable plasma viremia at necropsy and plasma viral RNA from viremic time points was sequenced using an Illumina-based sequencing approach as previously described [28] with primers SIV.INT.P5 (5ʹ-GAAGGGGAGGAATAGGGGATATG-3ʹ) and SIV.INT.P7 (5ʹ-CCTCCATGTGGGAACTGCTATCC-3ʹ). During a previous study, it was discovered that one viral lineage has a lower replication capacity [28] and was therefore removed from the analysis of the current manuscript. We renormalized the proportions of the viral load composed of the remaining barcodes for the analysis and figures in this current paper.

The experimental protocol for the intravenously inoculated animals was previously described in Khanal et al. [26] and Okoye et al. [27]. Briefly, 58 Indian-origin rhesus macaques were intravenously challenged with 100 (n=2), 200 (n=22), 500 (n=30), or 2200 IU (n=4) of SIVmac239M (∼10000 barcodes [25]). The two 100 IU and two 200 IU inoculated animals had plasma collected for viral RNA sequencing 8 and 14 dpi. The 500 IU and remaining twenty animals at the 200 IU dose, initiated ART 12 dpi, and plasma was collected 15 dpi for viral RNA sequencing. Finally, plasma was collected for viral RNA sequencing from the 2200 IU inoculated animals at 11 or 12 dpi (two animals each). Sequencing was again performed using the same Illumina-based sequencing approach previously described in Fennessey et al. [25] with primers VpxF1 (5’-CTAGGGGAAGGACATGGGGCAGG-3’) and VprR1 (5’-CCAGAACCTCCACTACCCATTCATC-3’).

#### In vitro cultures

Indian origin rhesus macaque peripheral blood mononuclear cells (PBMCs) were isolated from whole blood using SepMate tubes (StemCell Technologies) with Lymphoprep (StemCell Technologies) density gradient medium and centrifugation. CD4^+^ T cells were enriched by negative selection (CD4 T cell isolation kit; Miltenyi). Cells were activated with 5 μg/mL phytohemagglutinin for 3 days and cultured in RPMI supplemented with 10% fetal bovine serum (FBS), 2mM L-glutamine, 100 U/mL penicillin and 100 μg/mL streptomycin (RPMI-complete) supplemented with IL-2 (100 U/mL). Cells were infected with a stock of SIVmac239V67M that was pre-treated with DNase I (RQ1 RNase-Free DNase, Promega: 32 U/ml) to eliminate any residual plasmid DNA. Cells were infected at an MOI of 0.003 (as determined by TZM-bl infectivity assay). Infection was performed by spinoculation for 2 hours followed by overnight incubation at 37°C. After the overnight incubation (∼18 hours), cells were washed with pre-warmed PBS five times to minimize the presence of any residual DNA from the inoculum and resuspended in RPMI complete with IL-2 and treated with 10 μM maraviroc and 2 μg/mL anti-CD4 or left drug-free in culture. At 24 hours post infection, 1 μM emtricitabine (FTC) was added to the treated culture to block any additional rounds of replication. Supernatant was collected at pre-specified time points, with media replenished after each collection. SupT1-R5 cell line was similarly cultured in RPMI-complete and infected with the same SIVmac239V67M stock at an MOI of 0.005. Drugs were administered and samples collected as described above.

To quantify viral outgrowth and prepare samples for deep sequencing, cell-clarified supernatant was centrifuged at max speed (16,000 x *g*) for 1 hour to pellet virus. Viral RNA was then isolated using the Qiagen QIAamp Viral RNA mini kit. Complementary DNA (cDNA) was generated using SuperScript III (Invitrogen) and an SIV-specific reverse primer (SL8R: 5’ AGCTGAGAGAGGATTTCCTCCC 3’). cDNA was quantified via qRT-PCR using the primers VpxF1 5’-CTA GGG GAA GGA CAT GGG GCA GG-3’ and VprR1 5’-CCA GAA CCT CCA CTA CCC ATT CATC with a labeled probe (ACC TCC AGA AAA TGA AGG ACC ACA AAG GG). Prior to sequencing, PCR was performed with VpxF1 and VprR1 primers containing either the F5 or F7 Illumina adaptors with a unique 8-nucleotide index sequence for multiplexing. PCR was performed using High Fidelity Platinum Taq (ThermoFisher). The multiplexed samples were sequenced on a MiSeq instrument (Illumina) and analyzed as previously described [25, 26, 39].

The viral stock used in this study, SIVmac239V67M, is a molecularly barcoded synthetic swarm in which a 34-base insert containing a stretch of 10 random nucleotides was inserted between the vpx and vpr genes of the SIVmac239 clone containing a V67M amino acid modification in the envelope gene, as described previously [25]. The resultant barcoded SIVmac239V67M viral stock exhibits comparable replication rates to wild-type SIVmac239 in vitro (data not shown), contains ∼23,000 distinct barcode variants, and has an infectious titer of 2.8×10^5^ IU/mL.

### Mathematical modeling and statistical analysis

#### Quantification of lineage size heterogeneity for in vitro experiments with a large tail

As shown in the insets of Fig 2B and C, the distribution in lineage sizes in the in vitro experiments consisted of a lognormal distribution and a large tail of small lineages. Therefore, to calculate the variance of the lognormal component in each well, we fit a mixture distribution to the fraction of the viral load composed of individual lineages. For the tail component of this mixture distribution, we used a power-law distribution (also known as a Pareto distribution), which has a probability density function of the form

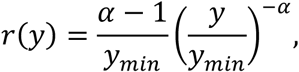

where α is the power of the distribution and *y_min_* is the minimum value of the variable *y*. Naturally, the remaining component of the mixture distribution was a lognormal distribution. Thus, if we define *x* as the fraction of the viral load composed of a given barcode, the full mixture distribution, *q(x)*, is given by

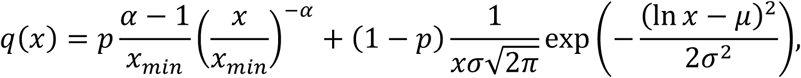

where *p* is the probability that a lineage is part of the power-law component as opposed to the log-normal component, *x_min_* is the minimum detected fraction of the viral load within the well, and μ and σ are the mean and standard deviation of the natural logarithm of *x* or lineages in the log-normal component of the distribution.

We employed a Maximum Likelihood approach to fit the mixture distribution to the data from each well.

The standard deviation can be converted from base *e* (σ) to base 10 (α_10_) via

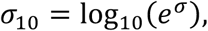

and therefore, the variance of log_10_ lineage size (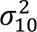; as reported in the Results) is given by

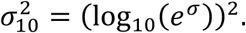

### Calculation of the proportion of lineages within 10-fold of geometric mean

To estimate the percentage of lineages within 10-fold of the geometric mean for each experimental set up (indexed *j*), we used the calculated lineage size heterogeneity (i.e., the average variance of log_10_ of the lineage sizes in set up *j* 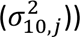.

We defined *s* to be lineage size normalized by the geometric mean lineage size, giving a corresponding log-normal cumulative distribution function, 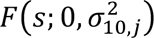. Note that we set the mean logarithm of *s* to be zero for this distribution function, because as *s* is normalized by the geometric mean lineage size, the mean of log_10_ *s* is 0 by definition. We then calculated the difference between the cumulative distribution function evaluated at distances of 10-fold away from the geometric mean, i.e. at *s* = 10 and *s* = 0.1,

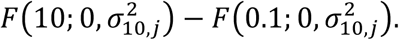

### Statistical comparison of lineage size variances

The statistical significance of differences in lineage size variance between experimental set ups was assessed using a standard two sample Student’s t-test (t.test from the stats R package).

### Estimation of proportional breakdown of dissemination bottleneck

We estimated the proportional contribution of sources of viral production heterogeneity (viral-mediated or phenotypic heterogeneity) to the total dissemination bottleneck by comparing lineage size variance caused by these sources of heterogeneity to the total variance in the most “disseminated” setup (i.e. intravenous inoculation). Prior to these comparisons, we first removed bias due to inherent variability in PCR amplification by background subtracting the lineage size variance in the PCR amplification data (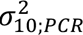 estimated from the low-input sequencing; S1 Text) from the mean observed variances for each experimental set up. For the purpose of simplicity for the remainder of this section we will refer to this background subtracted lineage size variance simply as lineage size variance, and label it as 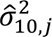, where

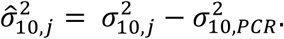

We estimated the proportional contribution of viral-mediated heterogeneity to the overall dissemination bottleneck as the ratio of (background subtracted) lineage size variance in the cell line experiments 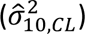 to that of the intravenously inoculated animals 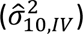, i.e.

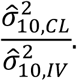

We estimated the proportional contribution of phenotypic heterogeneity to the overall dissemination bottleneck by considering the ratio of the increase in lineage size variance from the cell line to the stimulated primary cell experiments to the variance of the intravenously inoculated animals, i.e.

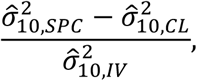

where 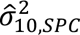 is the (background subtracted) variance in production by stimulated primary cells in vitro.

## Acknowledgments

None.

## Author contributions

Conceptualization: SSD, MPD, BFK

Methodology: SSD, AM, TES, CMF, DC, MPD

Formal analysis: SSD

Investigation: AM, CMF, BVM, LJP, AAO, TTI

Resources: BFK, LJP, MPD

Data Curation: SSD, BFK

Writing – Original Draft: SSD, MPD, DC, AM, CMF

Writing – Review & Editing: SSD, MPD, DC, BFK, AM, TES, CMF, BVM, LJP, AAO, TTI

## Supporting Information Captions

**S1 Fig. Correlation between number of detected barcodes and inoculation size**

**S1 Table. Quantitative summary of experimental setups**

**S1 Text. Assay and viral variability insufficient to explain lineage size heterogeneity**

**S2 Text. Lineage size distribution following untreated in vitro replication**

