## Supplementary material for "Dissecting the steps in early Simian Immunodeficiency Virus dissemination following mucosal and intravenous infection of rhesus macaques": S1 Fig

S1 Fig. Correlation between number of detected barcodes and inoculation size

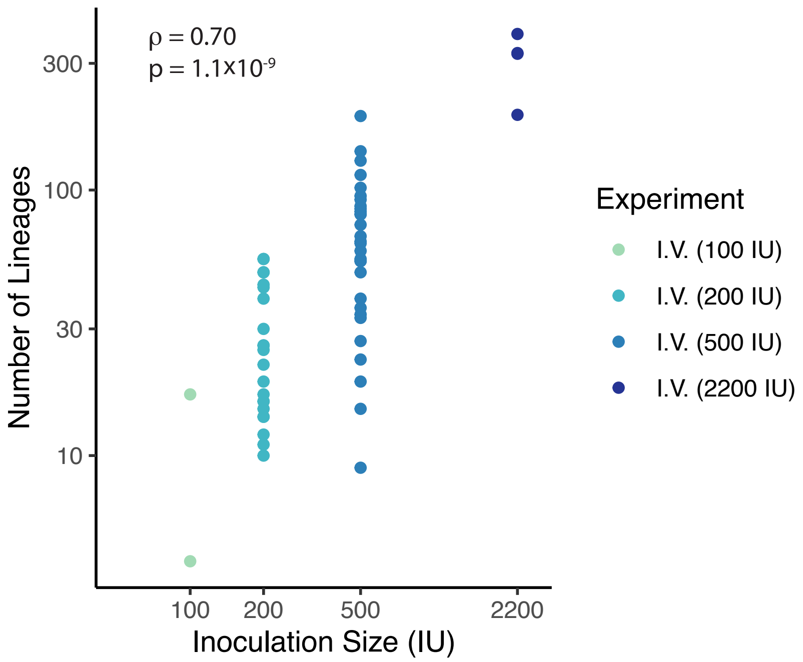


*Figure S1: Number of founding lineages is correlated with inoculation size. The number of unique barcodes detected within each intravenously inoculated animal plotted against the inoculation size used to infect the animal (reported in infection units (IU) as determined by TZM-bl infectivity assay). The Spearman correlation coefficient and corresponding p-value are listed.*
