## Supplementary material for "Dissecting the steps in early Simian Immunodeficiency Virus dissemination following mucosal and intravenous infection of rhesus macaques": S1 Table

S1 Table Quantitative summary of experimental setups

| Experimental Setup | Number of lineages detected (range) | Maximum fold-difference in size | Mean variance ($log_{10}$-scale)* | Mean variance prior to removal of tail ($log_{10}$-scale)*^ |
|---|---|---|---|---|
| Mucosal (intravaginal) inoculation | 2-8 | $1.09 \times 10^3$ | $1.24 \pm 0.54$ | |
| Intravenous (100 IU) inoculation | 4-17 | $1.73 \times 10^3$ | $0.75 \pm 0.25$ | |
| Intravenous (200 IU) inoculation | 10-55 | $2.08 \times 10^5$ | $1.21 \pm 0.12$ | |
| Intravenous (500 IU) inoculation | 9-190 | $3.76 \times 10^5$ | $1.46 \pm 0.09$ | |
| Intravenous (2200 IU) inoculation | 192-387 | $9.97 \times 10^4$ | $1.26 \pm 0.10$ | |
| Stimulated primary cells (treated) | 1192-1305 | $1.42 \times 10^3$ | $0.31 \pm 0.01$ | $0.82 \pm 0.03$ |
| SupT-R5 cell line (treated) | 2932-3023 | $8.69 \times 10^2$ | $0.17 \pm 0.002$ | $0.99 \pm 0.01$ |
| Low-input sequencing | 29-42 | 5.37 | $0.013 \pm 0.001$ | |
| Stimulated primary cells (untreated) | 664-772 | $1.55 \times 10^3$ | $0.28 \pm 0.01$ | $0.50 \pm 0.03$ |
| SupT-R5 cell line (untreated) | 1614-1692 | $8.54 \times 10^3$ | $0.45 \pm 0.02$ | $0.75 \pm 0.02$ |

*± standard error. ^Tail only removed from in vitro experiments.
