## Supplementary material for "Dissecting the steps in early Simian Immunodeficiency Virus dissemination following mucosal and intravenous infection of rhesus macaques": S1 Text

### S1 Text Assay and viral variability insufficient to explain lineage size heterogeneity

In this supplement, we confirm that the observed lineage size heterogeneity is genuine and not the result of measurement error or a unique characteristic of the experimental model used. Specifically, we demonstrate that the two alternative hypotheses suggested in the main text; (i) measurement error due to PCR amplification bias and (ii) differences in viral fitness among barcoded clonotypes, are insufficient to explain the observed lineage size heterogeneity.

#### S1.1 Assessment of the potential impact of PCR amplification bias

High variability in PCR amplification of lineages may have artificially increased the lineage size variance. If this was the case, then the size of individual lineages should not be correlated between samples taken from a single animal at different time points (since the sizes would be based on variability in the PCR amplification). We sequenced plasma virus from the 100 IU and two 200 IU intravenously inoculated animals six days after their first sequenced sample (day 8) and observed a strong correlation in lineage sizes between the two time points (Figure S1.1A; Pearson correlation coefficient of 0.95, $p < 10^{-24}$) indicating observed lineage size heterogeneity is not solely due to variability in PCR amplification.

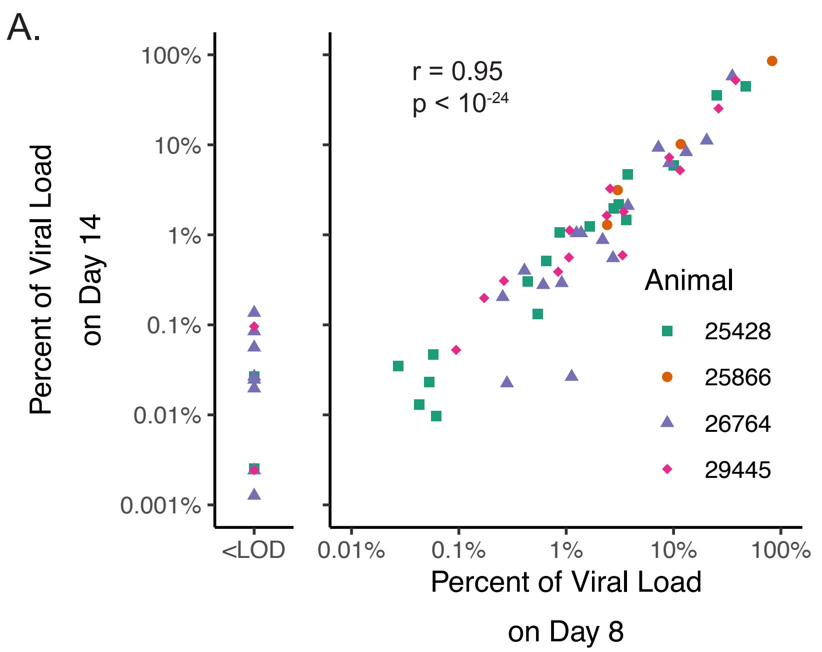


*Figure S1.1: PCR amplification bias is insufficient to explain lineage size heterogeneity. (A) Positive Pearson correlation in the percentage of viral load composed of individual barcoded lineages between plasma samplings on days 8 and 14 in four intravenously inoculated animals. Two animals were inoculated with 100 IU (25428 and 25866) and two with 200 IU (26764 and 29445). Lineages newly detected on day 14 are indicated as below limit of detection (LOD) on day 8 and are not included in calculation of correlation coefficient and significance. (B) Empirical distributions in PCR amplification of single viral templates from 1:9.8 million dilutions of SIVmac239M2 barcoded viral stock.*

In order to directly measure the variability in PCR amplification of individual viral templates we diluted the viral stock 9.8 million-fold, with the idea being that at this dilution we would obtain at most one template of a given barcode in a well. We considered 10 replicates and found that each replicate contained less than 0.03% of the stock barcodes (≤42 barcodes out of more than 140000 barcodes in the stock [1]), confirming that it was highly unlikely that any single barcode was present at more than one copy.

Following PCR amplification, the sequence count for individual barcodes had an average variance of $0.013 \pm 0.001$ ($\log_{10}$ copies)$^2$ (mean $\pm$ standard error) across the ten replicates (range 0.008 to 0.021; Figure S1.1B). This variability in amplification of individual templates is minor (1%) compared to the observed spread in lineage size in vivo, as illustrated in Fig 3A of the main text.

#### S1.2 Assessment of replication bias among barcoded viral clonotypes

Another potential cause of lineage size heterogeneity not reported on in the Results of the main text is the potential variation in replicative capacity across viral lineages. If different clonotypes had different replicative fitness in this experimental model, faster replicating clonotypes would tend to be larger and grow to be even more dominant in the plasma viral load as infection progresses. Additionally, we would expect more fit barcodes to be detected in a larger proportion of animals and consistently be the dominant barcodes within those animals.

The same four animals that were sequenced at two separate time points and analysed in S1.1 can also be used to assess variation in barcode growth rate. Comparison of increases in barcoded clonotype viral loads from day 8 to 14 in these animals demonstrates that barcoded lineages in fact grow in parallel (Figure S1.2), suggesting that variation in replicative fitness is negligible.

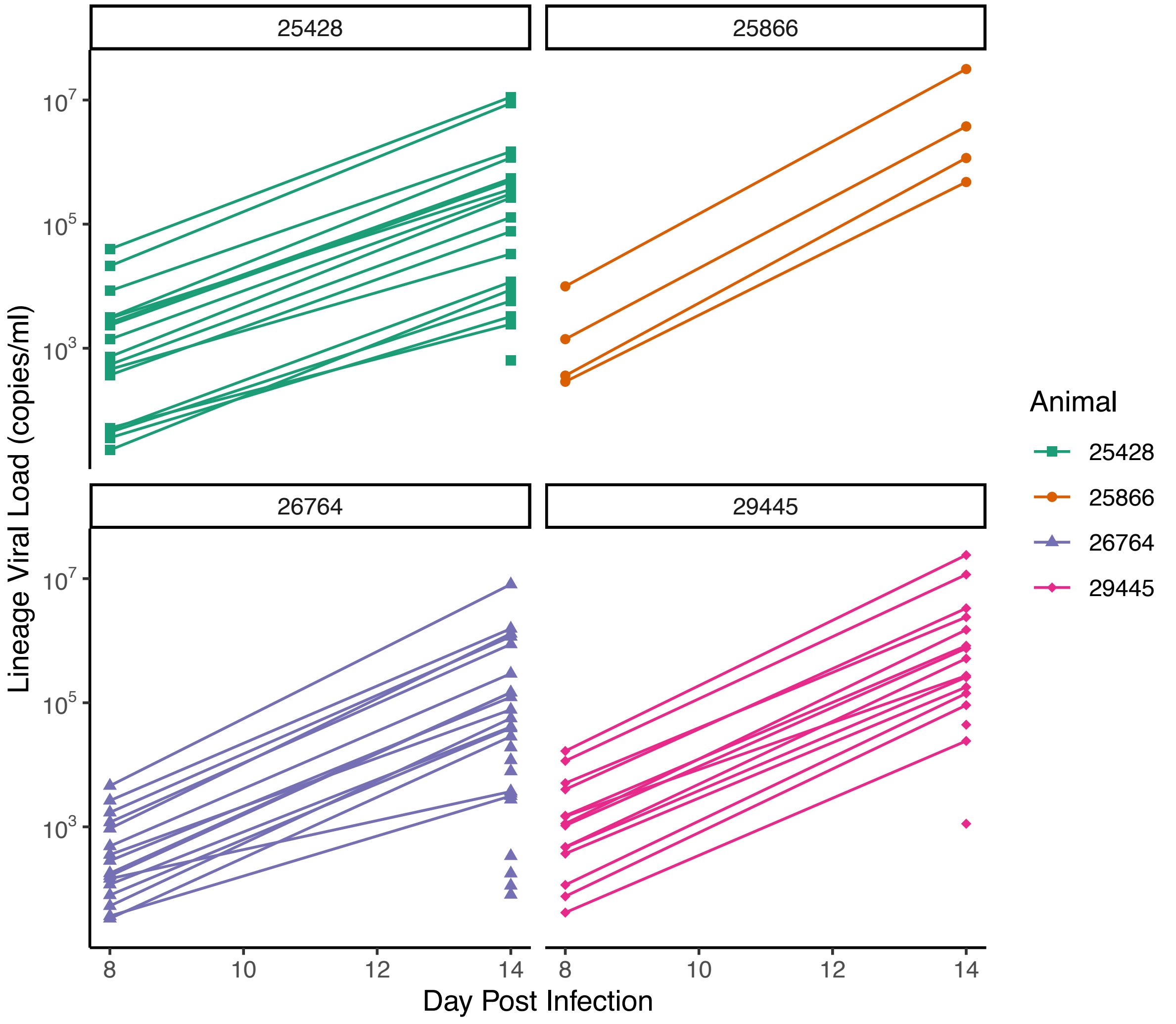


*Figure S1.2 Parallel growth of barcoded viral lineages in vivo. Viral load of individual barcoded lineages on 8 and 14 days post infection in four animals inoculated intravenously with 100 IU (25428 and 25866) or 200 IU (26764 or 29445). Most barcodes were detected at both time points, and lines connect viral loads on day 8 and 14 corresponding to the same barcode.*

Furthermore, we saw no evidence of select barcodes being overrepresented across animals. In fact, only a single barcode was the largest clonotype in more than one animal (Figure S1.3A) and 71.5% of all barcodes detected in an intravenously inoculated animal were detected in only one animal (Figure S1.3B).

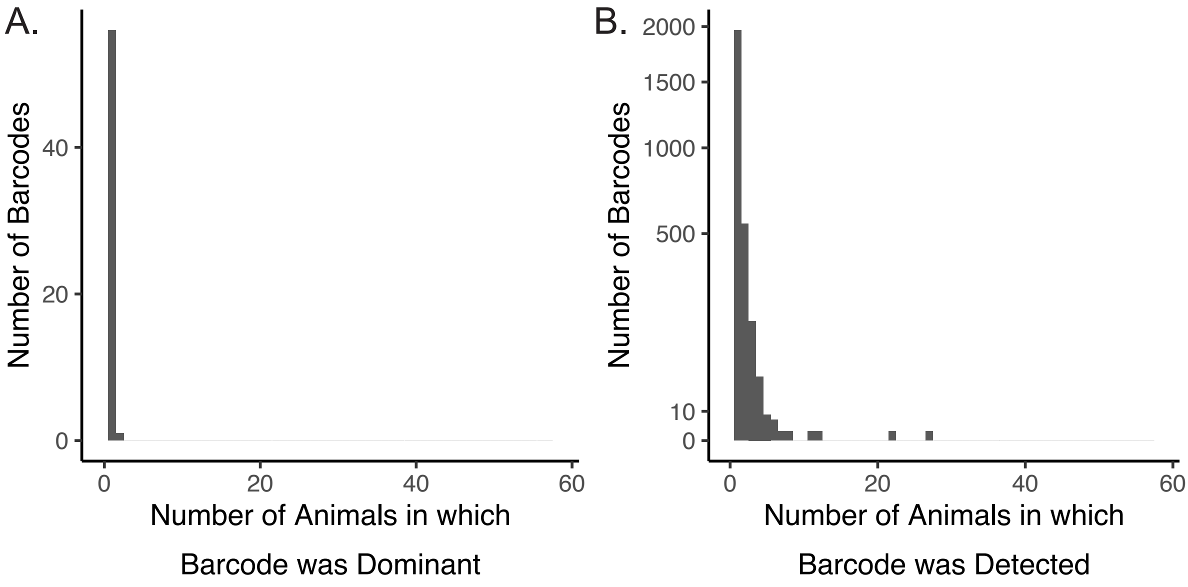


*Figure S1.3 Different barcodes detected in different animals. Histograms for (A) the number of animals in which a barcode was the largest barcode or (B) simply detected in the plasma viral load. Histograms were generated based on data from all animals intravenously inoculated with SIVmac239M (n = 58).*

Taken together, these data demonstrate that variation in viral fitness is not a driver of lineage size heterogeneity in this experimental model. However, in the context of exposure to a diverse viral pool, lineage size heterogeneity would most likely be exacerbated by discrepancies in replicative fitness.
