## Supplementary material for "Dissecting the steps in early Simian Immunodeficiency Virus dissemination following mucosal and intravenous infection of rhesus macaques": S2 Text

S2 Text Lineage size distribution following untreated in vitro replication

The heterogeneity in lineage size observed following multiple rounds of replication is illustrated in Figure S2.1.

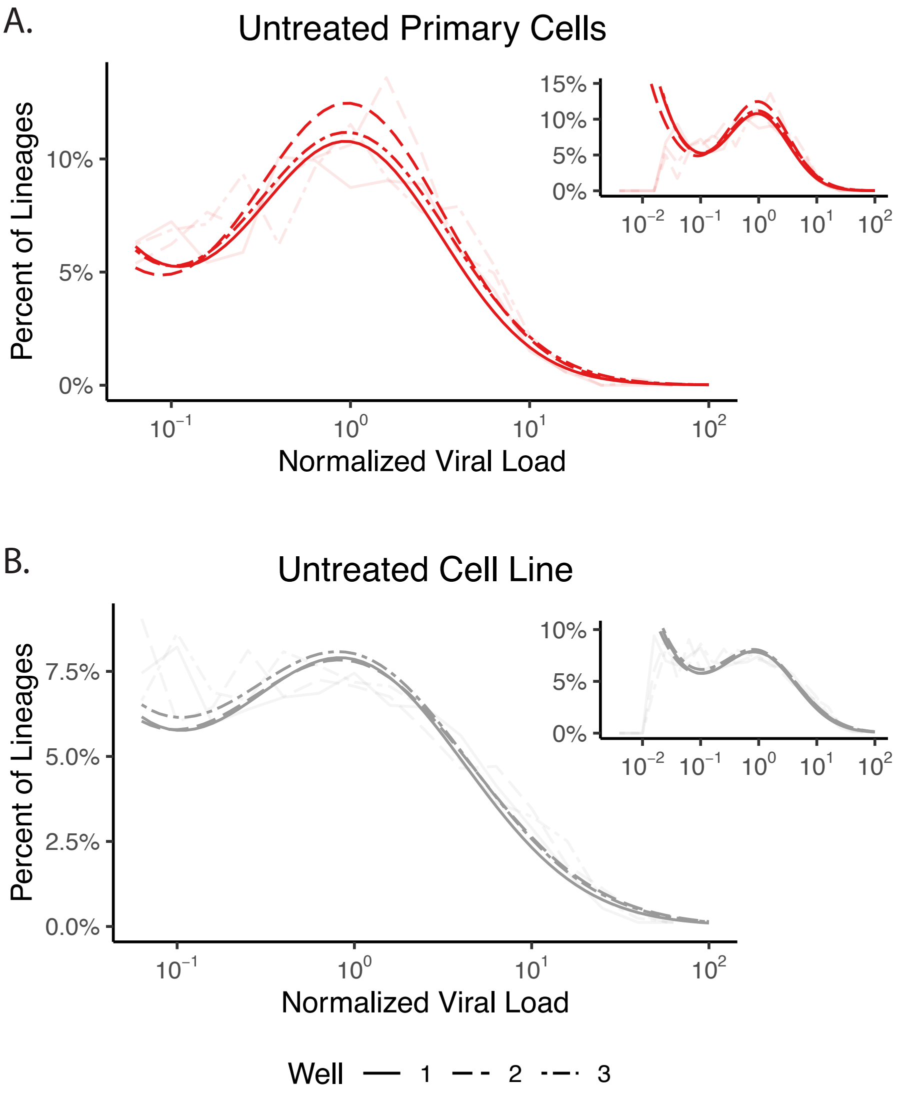


*Figure S2.1: Lineage size heterogeneity following untreated viral replication in vitro. Stimulated primary cells (A) or SupT1-R5 cells (B) were infected with SIVmac239V67M. Inoculum was washed out after ~18 hours, and supernatant was subsequently washed at 24 hours post infection and 24-hourly thereafter. Lineage sizes were measured via Illumina sequencing of the supernatant RNA at 4 days and 7 days after infection in the stimulated primary cells and SupT1-R5 cells, respectively. Empirical distributions (faded lines) and best fit distributions (dark lines) of lineage size ($log_{10}$-scale) in each of the three stimulated primary cell (A, red lines), and SupT1-R5 (B, grey lines) wells are plotted. Main figures focus on the log-normal component of the distributions, while insets display the full data sets and fits (including the tail).*

Here we explain how we estimated the potential contribution of multiple rounds of viral replication to the overall dissemination bottleneck. This estimate is based on comparison of lineage size diversity on day two of anti-retroviral treated in vitro wells (i.e., viral production by single cells; Fig 2 of the main text) to that on day four or seven of untreated wells (i.e., after multiple rounds of viral replication has occurred; Figure S2.1). Lineage size variance increased from single cell production to following multiple rounds of replication in SupT1-R5 cells but decreased in the stimulated primary cell experiments. Therefore, we use the increase in lineage size variance in the SupT1-R5 cell data to estimate an upper limit of the contribution of multiple rounds of viral replication to the overall dissemination bottleneck. Specifically, the difference in lineage size variance in treated and untreated experiments gives an estimate of the lineage size variance caused by multiple rounds of stochastic viral replication, and the ratio of this difference and the lineage size variance of the intravenously inoculated animals gives the estimated proportional contribution of multiple rounds of replication to the overall dissemination bottleneck.

We first removed bias due to inherent variability in PCR amplification by background subtracting the lineage size variance in the PCR amplification data ($\sigma^2_{10;PCR}$ estimated from the low-input sequencing; S1 Text) from the mean observed variances for each experimental set up (similar to comparisons in the main text). For the purpose of simplicity for the remainder of this Supplement, we will refer to this background subtracted lineage size variance simply as lineage size variance, and label it as $\hat{\sigma}^2_{10,j}$ for experimental set up $j$, where

$$\hat{\sigma}^2_{10,j} = \sigma^2_{10,j} - \sigma^2_{10,PCR}.$$

We then estimated the upper limit on the proportional contribution of multiple rounds of replication to the overall dissemination bottleneck by considering the ratio of the increase in lineage size variance from treated to untreated in vitro cell line experiments to the lineages size variance of the intravenously inoculated animals, i.e.

$$Estimated\ contribution\ of\ multiple\ rounds\ of\ replication = \frac{\hat{\sigma}^2_{10,CLU} - \hat{\sigma}^2_{10,CL}}{\hat{\sigma}^2_{10,IV}},$$

where $\hat{\sigma}^2_{10,CLU}$, $\hat{\sigma}^2_{10,CL}$, and $\hat{\sigma}^2_{10,IV}$ are the (background subtracted) variances in lineage size measured in the untreated cell line in vitro wells, treated cell line in vitro wells, and the intravenously inoculated animals, respectively.
